# Role of Nrp1 in controlling cortical interhemispheric circuits

**DOI:** 10.1101/2021.05.12.443798

**Authors:** F Martín-Fernández, C. G. Briz, M. Nieto

## Abstract

Callosal projections establish topographically organized maps between cortical areas. Neuropilin-1 (Nrp1) cortical gradient induces an early segregation of developing callosal axons. We investigated later roles of Nrp1 on the development of callosal projections from layer (L) 2/3 of the primary (S1) and secondary (S2) somatosensory (SS) areas, which express higher and lower levels of Nrp1, respectively. We used *in utero* electroporation to knock down or overexpress Nrp1 combined with retrograde tracers, to map connections at postnatal day 16 and 30. High levels of Nrp1 blocked contralateral S2 innervation while promoted the late postnatal growth of homotopic S1L2/3 and heterotopic S2L2/3 branches into S1. Conversely, knocking down Nrp1 increased the growth of heterotopic S1L2/3 projections into S2, and the overall refinement of S2L2/3 branches, thereby diminishing the number of P30 S2L2/3 callosally projecting neurons. Thus, the Nrp1 gradient determines homotopic SSL2/3 callosal connectivity by regulating late postnatal branching and refinement in a topographic manner.

## Introduction

The cerebral cortex is responsible for the execution of higher cognitive functions (Hill and Walsh, 2005). During evolution, the cortex had increased in size and complexity permitting eutherian brains to acquire the corpus callosum (CC). The CC is a tridimensional structure of myelinated interhemispheric axons that mediates the higher processing of information by establishing a topographically and hierarchically organized communication. It interconnects neurons located in equivalent areas of the hemispheres (homotopic callosal connections), as well as neurons of different modalities and orders (heterotopic connections) (Wise and Jones, 1976; Miller and Vogt, 1984; Rakic, 1988; Fenlon *et al.*, 2017; De León Reyes *et al.*, 2020).

For the correct processing of information, developmental mechanisms must ensure the precise definition of two aspects of the organization of adult CC circuits: firstly, the number of callosally projecting neurons (CPNs) in each cortical area and layer, and secondly, the topographical arrangement of their contralateral axons. Both of these aspects are the result of developmental selection processes including refinement (Aboitiz and Montiel, 2003; Fame *et al.*, 2011; Fenlon and Richards, 2015). Not all cortical areas contain the same number of CPNs. Their precise proportion defines the different functional cortical areas and regions. Associative areas, for instance, contain more CPNs than primary regions. The layer (L) distribution of CPNs also varies within areas, although in general, CPNs are more abundant in L2/3 and L5 of the adult cortex, there are some in L6 and very few in L4. The selection of the axonal targets of these CPNs is also highly specific. Callosal axons branch and synapse in topographically reproducible locations in the contralateral hemisphere. As a rule, they branch more profusely in homotopic areas and less in heterotopic locations. These branches form axonal columns that are usually in proximity to the border between areas (Mitchell and Macklis, 2005; Courchet *et al.*, 2013; Suarez *et al.*, 2014; Rodriguez-Tornos *et al.*, 2016; Fenlon *et al.*, 2017).

As mentioned, axonal refinement plays an important role in CC development. On the one hand, postnatal refinement dictates the elimination of exuberant branches that had invaded the cortical plate but do not establish synapses efficiently. This is thought to select optimal functional connectivity (Stanfield *et al.*, 1982; Innocenti and Clarke, 1984; Dehay *et al.*, 1986; Meissirel *et al.*, 1991; Innocenti, 2020). On the other hand, developmental refinement determines the number of CPNs in each area and layer. Early in development, CPNs are remarkably exuberant. Many cortical neurons, and virtually all neurons located in the upper layers (L2/3 and L4), develop transient callosal axons that invade the contralateral territories and bear the potential to establish a mature callosal connection. Most of these developmental projections do not progress into mature interhemispheric connections and are instead eliminated during the first postnatal weeks of the animal’s life (De Leon Reyes *et al.*, 2019). This CPN refinement is mediated by mechanisms that partly depend on activity, but which are largely unknown (Innocenti and Clarke, 1984; Koralek and Killackey, 1990; Innocenti and Price, 2005; Mizuno *et al.*, 2007; Huang *et al.*, 2013; Suárez *et al.*, 2014; Antón-Bolaños *et al.*, 2019; De Leon Reyes *et al.*, 2019).

In the mouse developing cortex, Neuropilin-1 (Nrp1) is expressed in a high to low mediolateral gradient (Zhao *et al.*, 2011; Zhou *et al.*, 2013; Muche *et al.*, 2015). Nrp1 null mutant mice are embryonically lethal (Kitsukawa *et al.*, 1997). To circumvent lethality, previous studies relied on Nrp1 conditional lines and on mutants of the binding domain of the Semaphorins, which are among its known ligands (Nrp1^Sema-^ mutant). Nrp1 plays several roles during CC development (Hatanaka *et al.*, 2009; Zhao *et al.*, 2011; Zhou *et al.*, 2013). At the CC midline, the interaction of Nrp1 with Semaphorin 3 (Sema3) C mediates the crossing of callosal axons (Gu *et al.*, 2003; Niquille *et al.*, 2009; Piper *et al.*, 2009; Mire *et al.*, 2018). Sema3A, another ligand of Nrp1, is expressed in the developing neocortex in a gradient that is opposite and complementary to that of Nrp1 (Tamamaki *et al.*, 2003; Zhao *et al.*, 2011). The binding of Sema3A to Nrp1-PlexinA1 induces axonal repulsion via the collapse of axonal growth cones (Takahashi *et al.*, 1999; Fournier *et al.*, 2000; Wu *et al.*, 2014). Since due to the gradients, callosal axons from motor areas express high Nrp1 and low Sema3A levels, axonal repulsion leads to the segregation of motor and somatosensory (SS) callosal axons, which express the opposite combination (Zhou et al., 2013). This determines that motor and SS axons occupy the dorsal and ventral callosal routes, respectively, and contributes to their guiding to motor and SS contralateral areas. Accordingly, genetic ablations of Sema3A or Nrp1 disrupt this axonal order and disorganizes axonal projections in the contralateral hemisphere (Zhou *et al.*, 2013). This effect is mediated by the steep gradient of Nrp1 expression established between motor and SS areas. However, little is known about the consequences of the lower differences that Nrp1 expression gradient creates within each area. Also, it is not known if Nrp1 plays additional functions on callosal development other than guidance. Herein we investigated the roles of Nrp1 gradient during the organization of somatosensory interhemispheric maps.

## Results

### Nrp1 expression levels determine the pattern of SS contralateral innervation

To investigate the roles of Nrp1 in the development of the callosal circuits of the SS cortex, we performed *in utero* electroporation (IUE) of constructs knocking down (shNrp1) or overexpressing Nrp1 (CAG-Nrp1). IUE was performed at embryonic day (E) 15.5 to specifically target L2/3 neurons. Vectors were co-electroporated with a plasmid encoding GFP (CAG-GFP), thus allowing to characterize the electroporated neurons and their projections at selected stages after birth (Figure 1A). Electroporations were targeted to the SS cortex, which is functionally divided into the primary somatosensory cortex (S1) and the secondary somatosensory cortex (S2), which receives first and higher-order sensory inputs from the thalamus, respectively (Rakic, 1988; Watson, 2012). For the analyses, the S1 barrel field area and the more lateral S2 area (Figure 1B) were distinguished by anatomical hallmarks such as the high density of L4 DAPI^+^ nuclei in the barrels (Paxinos and Franklin, 2004). We first examined the effects of our Nrp1 manipulations on the mature circuit of postnatal day (P) 30 animals. Coronal sections of brains electroporated with the control plasmid (CAG-GFP) showed that callosal projections from GFP^+^ L2/3 neurons reproducibly elaborate separated axonal columns in the SS areas of the contralateral hemisphere as described (Courchet *et al.*, 2013; Suárez *et al.*, 2014). The main column is located at the border of the S1 and S2 area, hereafter referred to as S1/S2 column (Figure 1A-C, blue arrowheads). Another less dense but very similar column forms in the lateral border of S2, hereafter referred to as the S2 column (Figure 1A-C, magenta arrowheads). Within the S1/S2 column, axons branch more profusely in L2/3 and L5, and within the S2 column, in L2/3 (Figure 1C). When knocking down Nrp1, we did not advert any obvious change in the overall pattern of contralateral innervation (Figure 1D). By contrast, overexpressing Nrp1 in L2/3 neurons caused a visible reduction of S2 axons together with an increase of S1 innervation (Figure 1E). To quantify these phenotypes, we measured the pixels occupied by the GFP fluorescence signal in specific regions of interest (ROI) delineating the main relevant SS areas and columns. To account for any differences in electroporation efficiency, the values of GFP within these ROIs were normalized to the fluorescence signal of the ipsilateral hemisphere (see Methods) (Rodriguez-Tornos *et al.*, 2016; Briz *et al.*, 2017). Firstly, the analysis of the total contralateral innervation showed that the average values and dispersion are indistinguishable in controls, shNrp1, and CAG-Nrp1 conditions (Figure 1F). This result confirmed that changing Nrp1 levels does not cause overall impairments of axonal innervation. Secondly, we quantified the number of GFP^+^ branches forming the S1/S2 column (Figure 1G), the S2 column (Figure 1H), and the axons in the remaining S1 area (Figure 1I). This quantification revealed a reduced S2 column and a slight increase of GFP^+^ axons in S1 in CAG-Nrp1 electroporated brains. No differences were detected in brains electroporated with shNrp1 (Figure 1G-I). An alternative analysis of the relative distribution of GFP^+^ axons in the different contralateral areas rendered equivalent results (See Methods and Figure 1 – figure supplement 1). The differences in axonal distribution were not due to neuronal death because the proportions of ipsilateral GFP^+^ neurons were indistinguishable in brains of all conditions (Figure 1 – figure supplement 2). Thus, increasing Nrp1 levels in L2/3 neurons blocks the development of their callosal axons in the contralateral S2 area while promoting branching in S1 territories. This suggests the implication of Nrp1 in the area-specific distribution of callosal branches.

**Figure 1.**
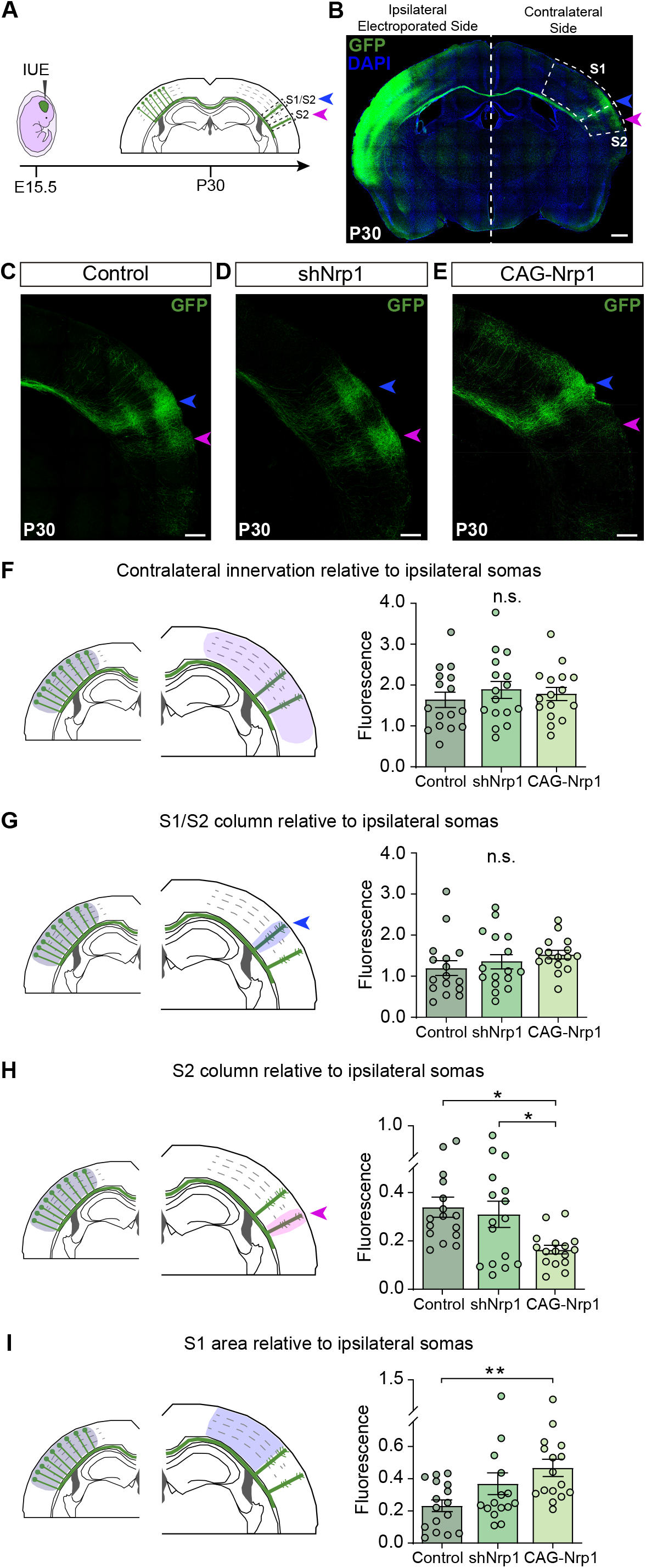
Analysis of the distribution of callosal axons upon alterations in Nrp1 levels. **A)** Scheme of the experimental approach. Contralaterally, L2/3 callosal axons form two main axonal columns, the S1/S2 column (blue arrow) and the S2 column (magenta arrow). **B)** Coronal section of P30 control brain electroporated at E15.5 with CAG-GFP. The SS cortex is divided into two functional areas: primary somatosensory cortex (S1) and secondary somatosensory cortex (S2) (dashed boxes). Green = GFP, Blue = DAPI. Scale bar = 500 μm. **C-E)** High magnifications of the contralateral hemisphere of P30 IUE brains showing GFP^+^ (green) axons of S1/S2 (blue arrow) and S2 columns (magenta arrow). Scale bar = 300 μm. **F-I)** Quantification of axonal distribution in the contralateral hemisphere. The left panels depict schemes showing the selected ROIs in which GFP^+^ is quantified (shaded areas). Graphs show values of GFP innervation relative to the fluorescence signal in the ipsilateral electroporated side of the same coronal section to normalize to the number of IUE L2/3 neurons. Mean ± SEM (n = 8 brains, 2 sections per brain, in all conditions). S1/S2 column (blue arrow), S2 column (magenta arrow). F) The innervation in the contralateral SS area. (One-way ANOVA: *P*-value = 0.6625 (n.s.)). G) S1/S2 column (One-way ANOVA: *P*-value = 0.3478 (n.s.)). H) S2 column (One-way ANOVA: *P*-value = 0.0085 (**). Posthoc with Tukey’s test: * *p*-value _Control – CAG-Nrp1_ = 0.0106, * *p*-value _shNrp1 – CAG-Nrp1_ = 0.0393). I) S1 area (One-way ANOVA: *P*-value = 0.0129 (*). Posthoc with Tukey’s test: ** *p*-value _Control – CAG-Nrp1_ = 0.0095).

### Nrp1 levels orchestrate callosal homotopic innervation in the somatosensory areas

We next analyzed the topographic location of the electroporated CPNs projecting to S1 or S2. Using stereotaxic coordinates, we performed classic axonal retrograde tracing by injecting fluorescent conjugates of cholera toxin subunit B of (CTB-555) in the cortical plate of the non-electroporated hemisphere. This procedure labels the subset of neurons projecting to the site of injection (Figure 2A). We injected P28 animals either in the S1 area, at the level of the S1/S2 column (Figure 2A-B), or in the S2 column (Figure 2A and C), and we analyzed the location of the GFP^+^CTB^+^ L2/3 CPNs at P30 (Figure 2A). As a retrospective control of the injection site, we confirmed that, in addition to cortical neurons, our injections in the S1/S2 column labeled preferentially thalamic neurons of the ventral posteromedial nuclei (VPM) (Figure 2 – figure supplement 1A-C), while our injections in the S2 column labeled neurons of the posterior nucleus (Po) (Figure 2 – figure supplement 1D-F) (see Methods). After counting the CPNs, we calculated the relative distribution of CPNs in S1 and S2 to evaluate changes in their contralateral targeting. For injections in the S1/S2 column, we calculated the ratio of GFP^+^CTB^+^ neurons in S1 vs. the number in S2 (homotopic projections vs. heterotopic projections). This analysis showed that in controls, most axons that form the S1/S2 column are homotopic projections from S1 since S1L2/3 CPNs were labeled 1.5 times more frequently than those in S2 (Figure 2D and J). Although there was a tendency to small decreases in the labeling of S1 CPNs, we observed no significant changes in shNrp1 or CAG-Nrp1 populations (Figure 2E-F and J). Hence, S1 innervation is not majorly affected by our manipulations. For animals injected in the S2 column, we calculated the ratio of GFP^+^CTB^+^ neurons found in S2 (homotopic) vs. those labeled in S1 (heterotopic) (Figure 2G-I, and K). This analysis showed that in controls, homotopic S2L2/3 projections are the main contributors to the GFP^+^ S2 column (2,5 ratio) (Figure 2G and K). Both knocking down or overexpressing Nrp1 decreased the proportions of S2L2/3 CPNs labeled with CTB injected in contralateral S2 (Figure 2H-I and K). The decreases observed in shNrp1 IUE brains indicated that their GFP^+^ S2 column is formed by an excess of heterotopic S1L2/3 axons (Figure 1D and H). Thus, the growth of shNrp1 S1L2/3 callosal branches compensates for the loss of Nrp1-deficient homotopic S2L2/3 projections. For the CAG-Nrp1 animals, since we had observed a reduction of the GFP^+^ S2 column (Figure 1E, and H), the data confirmed the loss of homotopic S2L2/3 branches. These shifts in the distributions of CPNs in shNrp1 and CAG-Nrp1 electroporated brains were not the consequence of differences in labeling efficiency, as we detected no changes in the distributions of non-electroporated CTB^+^ cells between conditions (Figure 2 – figure supplement 2). In sum, knocking down Nrp1 expression impairs the development of homotopic S2L2/3 callosal projections but is insufficient to trigger this effect in more medial S1L2/3 neurons. Contrary, incrementing Nrp1 expression blocks the development of both S1L2/3 and S2L2/3 branches into S2 equally, resulting in a diminished S2 column formed by an equilibrated proportion of S1L2/3 and S2L2/3 projections. These findings demonstrate that the gradient of Nrp1 expression favors homotopic SSL2/3 callosal connectivity.

**Figure 2.**
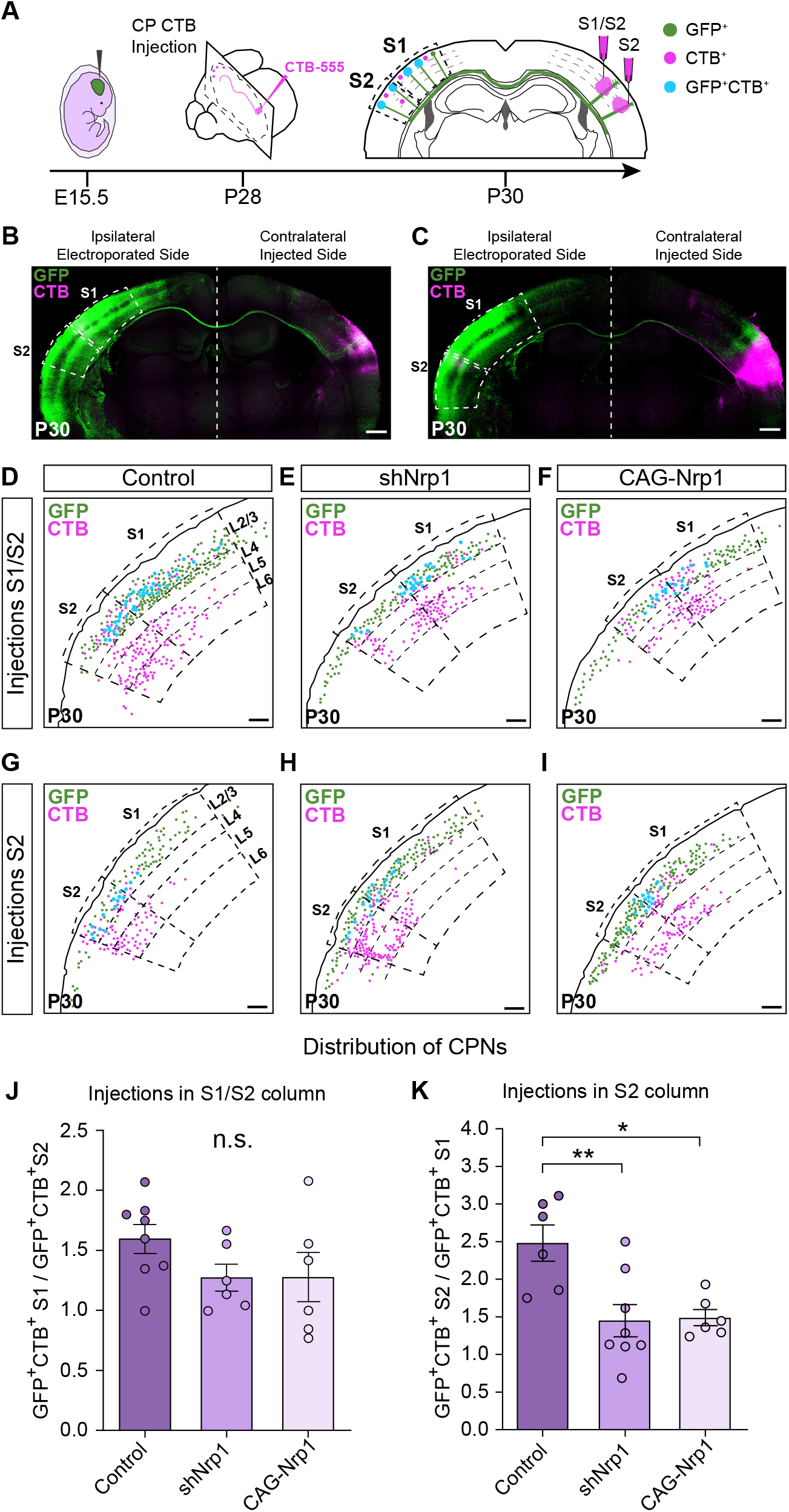
Analysis of homotopic and heterotopic projections in control, shNrp1, and CAG-Nrp1 IUE brains. **A)** Experimental workflow. After IUE at E15.5, brains are stereotaxically injected with CTB in CP at P28. Separate animals are injected in the S1/S2 column or the S2 column. Then, the numbers of GFP^+^CTB^+^ CPNs are quantified in S1 and S2 in the ipsilateral electroporated hemisphere at P30. **B-C)** Tilescan images of P30 coronal sections of control IUE brains injected in S1/S2 (B) or S2 coordinates (C). On the left, the ipsilateral side, where GFP^+^ and CTB^+^ somas are found. On the right, the contralateral hemisphere, the site of injections. Green = GFP, Magenta = CTB-555. Scale bar = 500 μm. **D-I)** Schemes reporting examples of the location of GFP^+^ neurons (green dots), CTB^+^ neurons (magenta dots), and GFP^+^CTB^+^ (blue dots) in coronal sections of IUE brains injected with CTB in the cortical plate. Scale bar = 300 μm. **J)** Quantification of the distribution of GFP^+^CTB^+^ after brains were injected in the S1/S2 column. Ratio (Y-axis) of the number of GFP^+^CTB^+^ in S1 divided by the number of GFP^+^CTB^+^ cells in S2 quantified in individual sections. Mean ± SEM (n ≥ 3 brains, 2 sections per brain in all conditions. One-way ANOVA: *P*-value = 0.2096 (n.s.)). **K)** Quantification of the distribution of GFP^+^CTB^+^ after brains were injected in S2. Ratio (Y-axis) of the number of GFP^+^CTB^+^ in S2 divided by the number of GFP^+^CTB^+^ cells in S1 quantified in individual sections. Mean ± SEM (n ≥ 3 brains, 2 sections per brain in all conditions. One-way ANOVA: *P*-value = 0.0036 (**). Posthoc with Tukey’s test: ** *p*-value _Control – shNrp1_ = 0.0052; * *p*-value _Control – CAG-Nrp1_ = 0.0109).

### Changes in Nrp1 expression alter developmental growth and refinement of callosal projections

Next, we investigated the mechanisms responsible for the different topography of callosal connectivity in Nrp1 electroporated P30 brains. To do so, we analyzed the innervation of P16 electroporated animals and compared them to P30. P16 control electroporated brains showed recognizable S1/S2 and S2 axonal columns, and quantifications demonstrated a distribution of contralateral branches very similar to that of P30 animals (Figure 3A, D-G). In contrast, P16 brains electroporated with shNrp1 or CAG-Nrp1 constructs showed reduced branching when compared to either P16 or P30 controls (Figure 3A-G). These reductions were observed throughout all contralateral areas (Figure 3 – figure supplement 1), although they were higher in S2 (Figure 3F). Besides, in controls, contralateral branches decreased from P16 to P30 (Figure 3D-F) as a consequence of the final pruning of callosal connections (O’Leary, 1992; De Leon Reyes *et al.*, 2019), but in shNrp1 and CAG-Nrp1 electroporated brains, axonal growth followed the opposite trend. Contralateral branches increased in all areas from P16 to P30, except in the S2 of CAG-Nrp1 brains (Figure 3D-G). Thus, abnormally late postnatal axonal growth compensates for the delayed contralateral innervation detected at P16, although with an incorrect topographic organization as revealed by our CTB labeling of CPNs.

**Figure 3.**
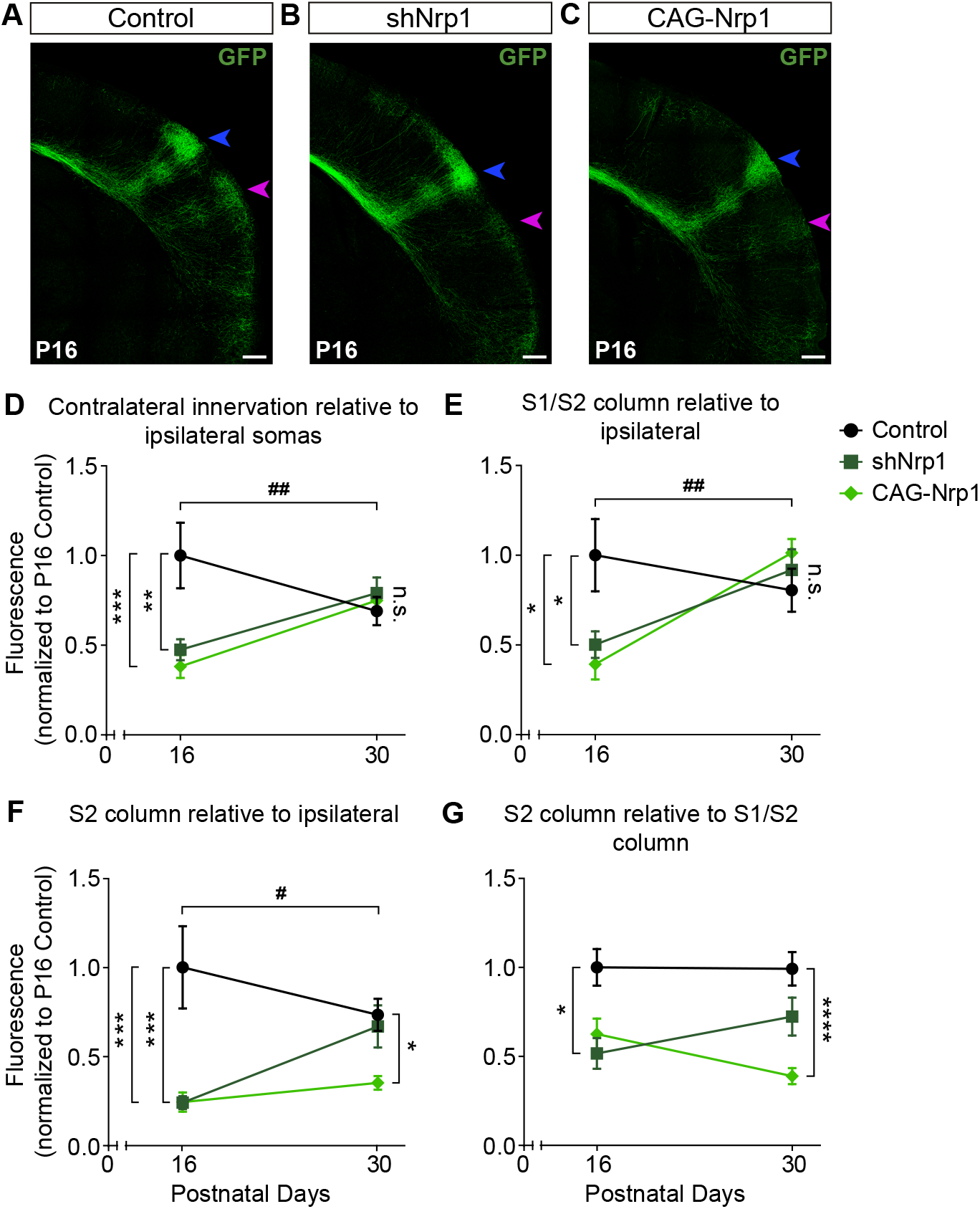
Comparisons of the postnatal changes of contralateral axons during the P16 to P30 window upon manipulations in Nrp1 expression. **A-C)** High magnification tilescan images of the contralateral hemisphere of IUE brains analyzed at P16 (Blue arrow = S1/S2 column. Magenta arrow = S2 column. Green = GFP). Scale bar = 300 μm. **D-G)** Plots of ratios of GFP^+^ innervation in the indicated area, relative to ipsilateral IUE hemisphere and normalized to the value of P16 control. Mean ± SEM (n ≥ 4 brains, 2 sections per brain in all conditions). D) Contralateral innervation in all SS area (Two-way ANOVA: *P*-value _Dynamics of contralateral innervation_ = 0.0012 (##); *P*-value _Postnatal day_ = 0.1224; *P*-value _Experimental condition_ = 0.0149. Posthoc with Tukey’s test: ** *p*-value _Control P16 – shNrp1 P16_ = 0.0044; *** *p*-value _Control P16 – CAG-Nrp1 P16_ = 0.0007). E) S1/S2 column (Two-way ANOVA: *P*-value _Dynamics of S1/S2 column_ = 0.0052 (##); *P*-value _Postnatal day_ = 0.0080; *P*-value _Experimental condition_ = 0.2043. Posthoc with Tukey’s test: * *p*-value _Control P16 – shNrp1 P16_ = 0.0465; * *p*-value _Control P16 – CAG-Nrp1 P16_ = 0.0116). F) S2 column (Two-way ANOVA: *P*-value _Dynamics of S2 column_ = 0.0114 (#); *P*-value _Postnatal day_ = 0.3331; *P*-value _Experimental condition_ < 0.0001. Posthoc with Tukey’s test: *** *p*-value _Control P16 – shNrp1 P16_ = 0.0003; *** *p*-value _Control P16 – CAG-Nrp1 P16_ = 0.0003; * *p*-value _Control P30 – CAG-Nrp1 P30_ = 0.0123). G) S2 column relative to S1/S2 column (Two-way ANOVA: *P*-value _Dynamics S2 column relative to S1/S2 column_ = 0.2098 (n.s.); *P*-value _Postnatal day_ = 0.8770; *P*-value _Experimental condition_ < 0.0001. Posthoc with Tukey’s test: * *p*-value _Control P16 – shNrp1 P16_ = 0.0102; **** *p*-value _Control P30 – CAG-Nrp1 P30_ < 0.0001). Data for P30 are from Figure 1 and Figure 1 – Figure supplement 1.

### Nrp1 levels determine the postnatal refinement of somatosensory L2/3 callosal axons at the midline

The CC organizes following dorsal-ventral topography. Decreasing Nrp1 expression in cortical motor neurons shifts callosal axon navigation to the ventral routes used by SS projections (Zhou *et al.*, 2013). Therefore, we analyzed the distribution of electroporated axons at the CC midline by quantifying the signal of GFP^+^ axons and plotting it against the dorso-ventral length of the CC divided into ten equal bins (Figure 4A). These analyses were performed at P16 and P30. In control, shNrp1 and CAG-Nrp1 electroporated brains, SS callosal projections crossed the midline by the most ventral two-thirds parts of the CC (bins 1-7) at both developmental stages. Only a few axons navigated by the most dorsal pathway (bins 7-10) (Figure 4B-I). Control, shNrp1, and CAG-Nrp1 P16 distributions were very similar, showing small differences in bins 3 and 5 (Figure 4B-E). Differences increased at P30. GFP^+^ callosal axons occupied slightly more dorsal routes (bins 4-10) in shNrp1 and CAG-Nrp1 electroporated brains than in controls (Figure 4F-I). We then compared, for each condition, the distributions at P16 and P30. This analysis revealed developmental changes. It showed that in controls, callosal axons that cross through the most dorsal paths (bins 3-8) are refined from P16 to P30, which suggests a topographic developmental elimination of interhemispheric projections (Figure 4B, F and J). This elimination of midline axons was reduced in shNrp1 electroporated brains, especially of those using dorsal routes (Figure 4K). Remodeling was not detected in CAG-Nrp1 electroporated brains (Figure 4L). The results suggested that changes in Nrp1 levels modify the developmental elimination of topographically organized callosal connections.

**Figure 4.**
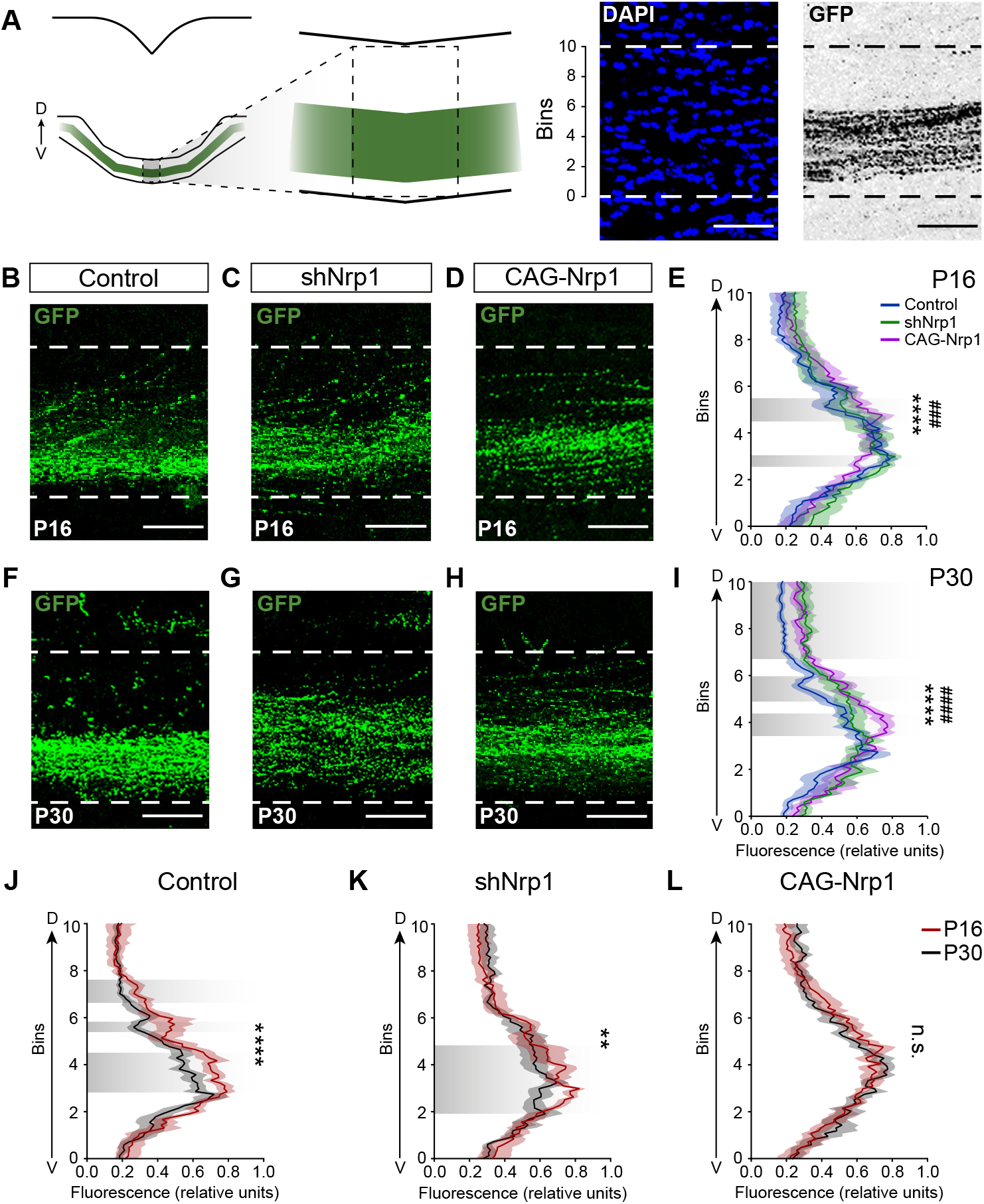
Analysis of the dorsoventral distribution of axons at the midline. **A)** Scheme of the analysis. The left panel depicts the selected ROI. The right panel shows the ROI divided into ten equal bins and applied to an image of the midline. DAPI image (blue) and pixels occupied by the GFP axons (grey). Scale bar = 50 μm. **B-D)** Images of the CC at the midline in P16 brains. (Green = GFP). Scale bar = 50 μm. **E)** Quantification of the dorsoventral distribution of GFP signal at P16 (ventral position, bins 0; dorsal position, bins 10). Lines represents mean ± SEM (shade) (n ≥ 3 brains, 2 sections per brain in all conditions). Shaded areas in grey indicate statistically significant differences (Two-way ANOVA: *P*-value _CC distribution_ = 0.9992; *P*-value _Bins_ < 0.0001; *P*-value _Experimental condition_ < 0.0001. Posthoc with Tukey’s test: **** *p*-value _Control - shNrp1_ 0.0001; ### *p*-value _Control – CAG-Nrp1_ = 0.0006). **F-H)** Images of the CC at the midline in P30 brains (Green = GFP). Scale bar = 50 μm. **I)** Quantification of the dorsoventral distribution of GFP signal at P30 (ventral position, bins 0; dorsal position, bins 10). Lines represents mean ± SEM (shade) (n ≥ 3 brains, 2 sections per brain in all conditions). Shaded areas in grey indicate bins showing statistically significant differences (Two-way ANOVA: *P*-value _CC distribution_ = 0.9925; *P*-value _Bins_ < 0.0001; *P*-value _Experimental condition_ < 0.0001. Posthoc with Tukey’s test: **** *p*-value _Control – shNrp1_ < 0.0001; #### *p*-value _Control – CAG-Nrp1_ < 0.0001). **J-L)** Comparison of the distributions of axons at the CC in P16 and P30 brains. Lines represents mean ± SEM (shade) (n ≥ 3 brains, 2 sections per brain in all conditions). J) Control (Two-way ANOVA: *P*-value < 0.0001 (****)). K) shNrp1 (Two-way ANOVA: *P*-value = 0.0021 (**)). L) CAG-Nrp1 (Two-way ANOVA: *P*-value = 0.5513 (n.s.)).

### Knocking down Nrp1 eliminates populations of S2L2/3 CPNs by refinement

Our results suggested that Nrp1 influences refinement. We recently showed that as part of their normal differentiation, L2/3 neurons first develop axons that project callosally, and subsequently, postnatal activity-dependent refinement eliminates most of these developmental callosal axons in area-specific manners, thereby selecting the populations of adult CPNs. The most intensive refinement of L2/3 CPNs occurs during the first two weeks of postnatal life (P1-P16). From then on, the proportion of L2/3 CPNs is very similar to that of the adult (De Leon Reyes *et al.*, 2019). To investigate a possible influence of Nrp1 in developmental CPN refinement, we set to analyze CPN numbers in the SS cortex of control and Nrp1 electroporated brains at P16 and P30. To this end, instead of in the cortical plate, we injected CTB-555 directly in the CC in the hemisphere opposite to the electroporation (Figure 5A-C). This procedure labels all neurons with an axon crossing the midline, including those in the process of developing or refining their callosal projections (De Leon Reyes *et al.*, 2019). In controls, after the injections, quantifications showed proportions of S1L2/3 and S2L2/3 CPNs undistinguishable to those previously reported in P16 and P30 WT mice, indicating that IUE does not alter CPN development (Figure 5 – figure supplement 1–2) (Fame *et al.*, 2011; De Leon Reyes *et al.*, 2019). Refinement of L2/3 CPNs, in both S1 and S2 areas, was not altered upon overexpression of Nrp1. However, electroporation of shNrp1 modified refinement (Figure 5 D-I). In P16 control brains, 50% of GFP^+^ S1L2/3 neurons were CTB^+^ (CPNs) (Figure 5D and E). This number increased to 65% in shNrp1-targeted S1L2/3 (Figure 5D and F), indicating that low Nrp1 expression delays axonal refinement. In S2, 40% of GFP^+^ L2/3 cells were CTB^+^ in controls or shNrp1 P16 electroporated brains (Figure 5G). Thus, since the numbers of P16 L2/3CPNs in these brains are equal or higher than in controls, the reduced GFP^+^ innervation that we observed in shNrp1 or CAG-Nrp1 electroporated brains is due to the scarce branching of GFP^+^ L2/3 axons in the contralateral cortical plate. At P30, 45% of GFP^+^ S1L2/3 and around 36% of GFP^+^ S2L2/3 control neurons were CPNs (Figure 5D and G). In shNrp1 P30 brains, the percentage of GFP^+^ S1L2/3 CPNs was indistinguishable from controls (Figure 5D) but the proportion of S2L2/3 CPNs was significantly reduced (Figure 5G-I). Thus, on the one hand, late postnatal refinement normalizes the transient increases of P16 S1L2/3 CPNs induced by knocking down Nrp1. On the other hand, refinement causes the exceeding elimination of S2L2/3 CPNs resulting in a significant decrease of their final numbers in the mature P30 circuit. Together, the data shows that by regulating axonal growth and refinement, Nrp1 levels determine homotopic callosal connectivity.

**Figure 5:**
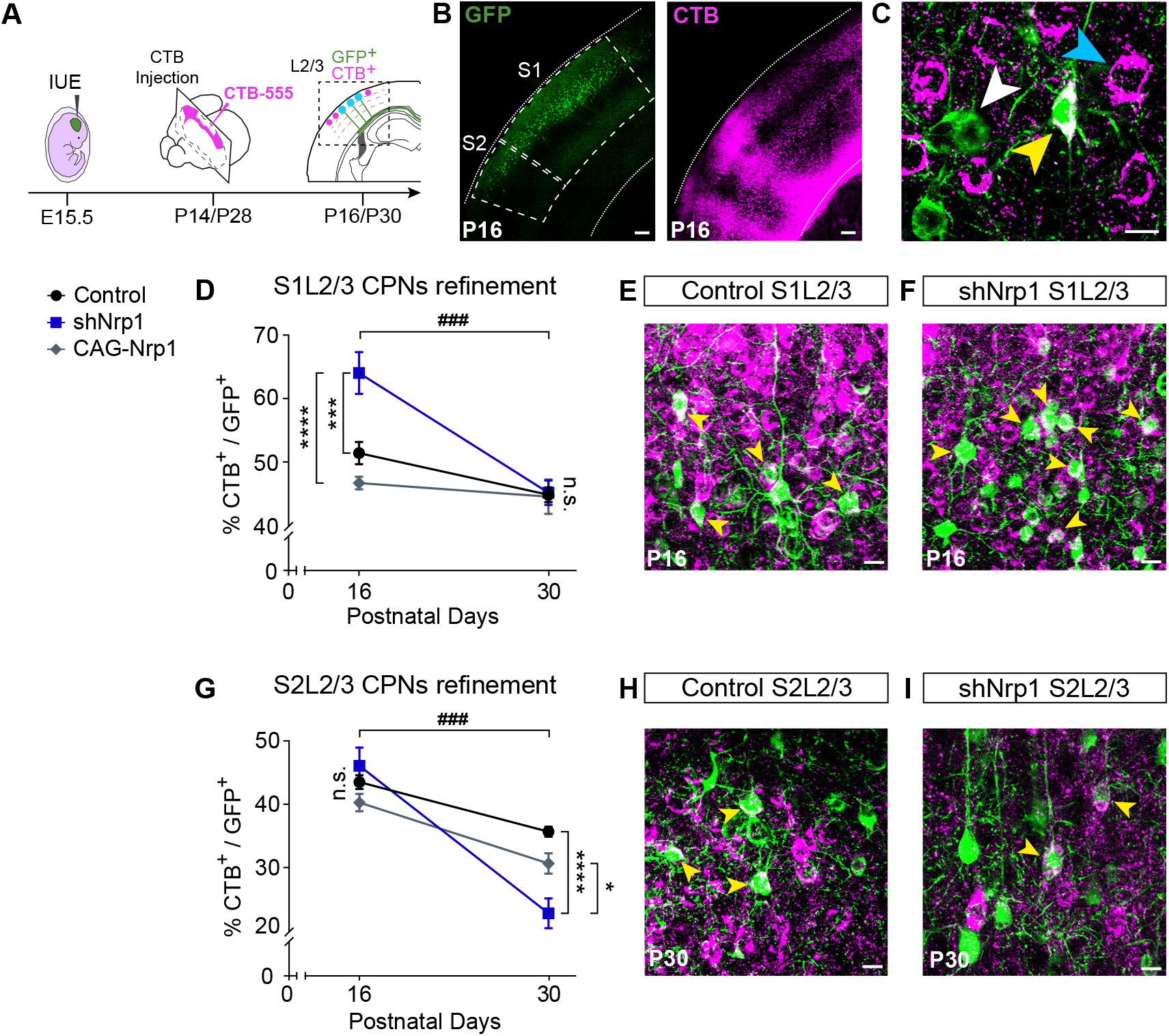
CPNs refinement during development (P16 to P30). **A)** Scheme of the experimental workflow. To analyze the effect of developmental refinement on the number of electroporated CPNs, stereotaxic CTB injection at the midline was performed after IUE at E15.5. **B)** Images showing ipsilateral cortex of P16 brains (delimitated by dashed lines). Left panel shows GFP^+^ electroporated neurons. Right panel shows the signal of CTB labeling axonal columns and somas (Green = GFP. Magenta = CTB-555). Scale bar = 300 μm. **C)** High magnification image of GFP^+^ L2/3 neurons in an injected P16 brain (GFP^+^, white arrowhead), (CTB^+^, blue arrowhead), (GFP^+^CTB^+^, yellow arrowhead). Scale bar = 10 μm. **D)** Plot of the proportion of CPNs (GFP^+^CTB^+^/GFP^+^) in S1 area at P16 and P30. Mean ± SEM (n ≥ 3 brains, 2 sections per brain in all conditions). (Two-way ANOVA: *P*-value _S1L2/3 CPNs refinement_ = 0.0007 (###); *P*-value _Postnatal day_ < 0.0001; *P*-value _Experimental condition_ = 0.0003. Posthoc with Tukey’s test: *** *p*-value _Control P16 – shNrp1 P16_ = 0.0003; **** *p*-value _shNrp1 P16 – CAG-Nrp1 P16_ < 0.0001). **E-F)** Merge images of control (E) and shNrp1 (F) S1L2/3 neurons at P16 (GFP^+^CTB^+^, yellow arrowheads). Scale bar = 10 μm. **G)** Plot of the proportion of CPNs (GFP^+^CTB^+^/GFP^+^) in S1 area at P16 and P30. Mean ± SEM (n 3 brains, 2 sections per brain in all conditions). (Two-way ANOVA: *P*-value _S2L2/3 CPNs refinement_ = 0.0003 (###); *P*-value _Postnatal day_ < 0.0001; *P*-value _Experimental condition_ = 0.0199. Posthoc with Tukey’s test: **** *p*-value _Control P30 – shNrp1 P30_ < 0.0001; * *p*-value _shNrp1 P30 – CAG-Nrp1 P30_ = 0.0127). **H-I)** Merge images of control (E) and shNrp1 (F) S1L2/3 neurons at P16 (GFP^+^CTB^+^, yellow arrowheads). Scale bar = 10 μm.

## Discussion

We herein demonstrate that, in the SS cortex, the Nrp1 gradient determines the topographic organization of SSL2/3 callosal connections. Nrp1 promotes homotopic branching and hinders heterotopic innervation (Figure 6). Previous studies have shown that the Nrp1 gradient in the cortex regulates an orderly organization of motor and SS axons during early postnatal development (Zhou *et al.*, 2013). We find that in addition to this function, Nrp1 regulates the late growth, branching, and terminal refinement of callosal axons. S1 and S2 areas process distinct somatosensory information received from first-order and higher-order thalamic nuclei (Inan and Crair, 2007; Pouchelon *et al.*, 2014). Thus, Nrp1 mediates a hierarchical organization of the bilateral exchange of sensory inputs. Our findings highlight the complex regulation required for the wiring of interhemispheric cortical maps.

**Figure 6.**
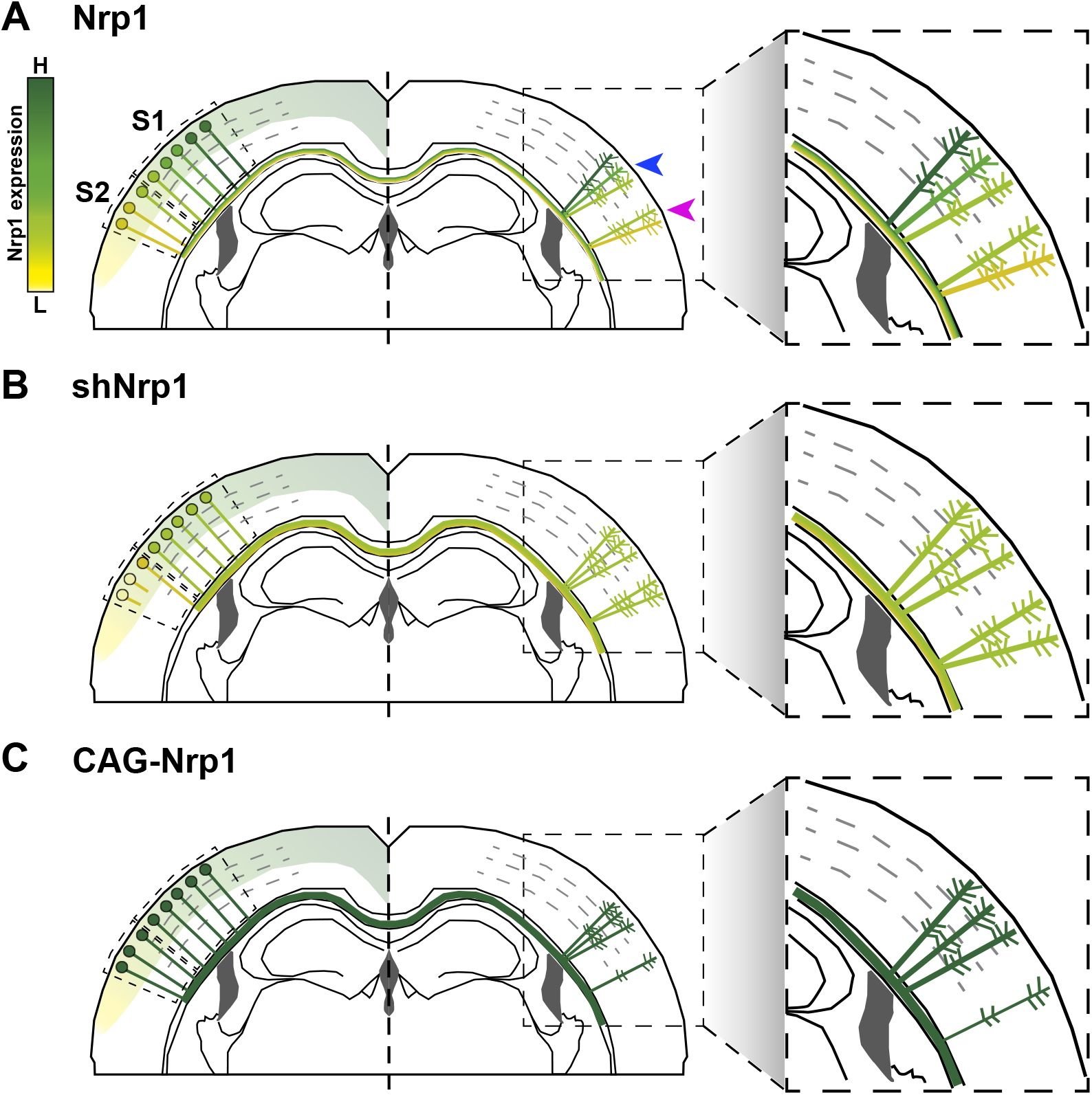
Model of the effects of Nrp1 levels on callosal connectivity. **A)** Nrp1 is expressed in the cortex in a high-medial, to low-lateral, gradient (higher levels in green, lower levels in yellow). Accordingly, S1L2/3 neurons (green dots) express high to intermediate levels of Nrp1. S2L2/3 neurons express low levels (light green and yellow dots). L2/3 neurons expressing high to intermediate levels of Nrp1 branch preferentially in homotopic S1/S2 column (blue arrow), and S2L2/3 neurons expressing intermediate to low levels into S2 areas (magenta arrow). **B)** Knocking down Nrp1 reduces the levels of Nrp1 according to the gradient. L2/3 CPNs with intermediate levels of Nrp1 expression can branch both in S1/S2 and S2. As consequence, an exceeding number of heterotopic branches from S1L2/3 CPNs outgrowth ectopically in the S2 column. They may outcompete axons of shNrp1 targeted S2L2/3 neurons that express very low levels of Nrp1 (light yellow dots). Many of these S2L2/3 neurons cannot terminate innervation and refine their callosal axon during the late period of P16-P30 developmental CPN refinement, thus becoming ipsilateral-only connecting neurons. **C)** Neurons over-expressing Nrp1 branch in the S1/S2 column but are not competent to innervate S2 areas, which is then significantly reduced.

We demonstrate that Nrp1 functions regulate the late development of callosal axonal branches. Our results argue in favor of possible mechanisms of competition or axonal cooperation, which are poorly studied in the CC (De León Reyes *et al.*, 2020; Innocenti, 2020). They cannot be merely explained by selective repulsion from the cortical plate, although they do not discard its contribution. For instance, we observe that knocking down or overexpressing Nrp1 delays the branching of SSL2/3 callosal axons in the cortical plate of P16 animals. These reductions are similar in all areas and both conditions, thus indicating that axons are not simply following a gradient. Instead, these P16 defects may reflect unbalanced ratios of stabilization/elimination of the synapses of callosal axons with their targets, which would decrease the rate of productive axonal branching and slow, but not block, cortical innervation (Courchet *et al.*, 2013). In agreement, shNrp1 and CAG-Nrp1 GFP^+^ callosal axons had innervated contralateral areas by P30. Competition is also suggested by the reconstitution of the S2 column in P30 shNrp1 IUE brains, which indicates that electroporated S1L2/3 branches outcompete homotopic S2L2/3 projections in these brains (Figure 6B). By contrast, the reduced P30 GFP^+^ S2 column in brains overexpressing Nrp1 is accounted for by the shift of all callosal projections towards the more medial S1 (Figure 6C). This is not in disagreement with repulsions, nor with other possible mechanisms. In every case, to dissect the connectivity produced by our manipulations of Nrp1, it is useful to examine why our manipulations alter the development of S2L2/3 callosal projections more than S1L2/3 CPNs. This is likely due to the nature of the endogenous gradient. While overexpressing vectors maximize the levels of Nrp1 equally in all neurons, the effects of shRNA constructs depend on the endogenous expression of the targeted transcripts and may result in intermediate and low Nrp1 levels in S1L2/3 and S2L2/3 neurons, respectively (Figure 6B). Accordingly, shNrp1-targeted S1L2/3 CPNs branching in S2 mimic the behavior of S2L2/3 WT neurons. Unfortunately, we could not assess the levels of Nrp1 protein in targeted electroporated neurons. In our attempts, antibody staining of Nrp1 did not detect the protein in the neuronal somas but only in the midline, similarly to other reports (Piper *et al.*, 2009; Zhao *et al.*, 2011; Zhou *et al.*, 2013; Lim *et al.*, 2015).

Hence, the effects of manipulating Nrp1 expression levels agree with that in the canonical WT circuit, interhemispheric axons from S1 neurons, which express higher levels of Nrp1, branch more profusely in homotopic S1 areas. Likewise, those projections from S2 CPNs, expressing lower Nrp1 levels, preferentially connect with contralateral homotopic targets in S2 (Figure 6A) (Yorke and Caviness, 1975; Wise and Jones, 1976; De León Reyes *et al.*, 2020). Electroporation of shNrp1 diminished homotopic S2L2/3 projections, which for a fraction of S2L2/3 CPNs, lead to the elimination of their callosal axons. These neurons presumably become ipsilateral-only projecting neurons, as it occurs to WT S1L4 and most other L2/3 cortical neurons during developmental normal refinement (Innocenti and Clarke, 1984; O’Leary and Koester, 1993; De Leon Reyes *et al.*, 2019). Interestingly, the refinement of these CPNs indicates that a certain level of Nrp1 expression is required for terminal callosal innervation. This again that not support axonal repulsion, may suggest a disadvantage at a possible axonal competition.

Sema3A seems a likely candidate responsible for the late branching phenotypes mediated by Nrp1. It has a developmental expression gradient opposite to Nrp1. This complementary expression determines repulsion (Kitsukawa *et al.*, 1997; Zhao *et al.*, 2011; Zhou *et al.*, 2013). In either case, a possible sequential role of Nrp1 in guidance and refinement is in agreement with observations in the cerebellum, where Nrp1 also has a dual function (Telley *et al.*, 2016). First, it guides inhibitory axons to their excitatory neuronal targets, and then, it determines the formation of synapses at specific locations within the neuronal body. The loss of Nrp1 in presynaptic basket cells blocks the formation of axonal synapses with Purkinje neurons, which resembles the inability of those Nrp1 deficient S2L2/3 CPNs to stabilize their callosal projections. In line with the involvement ofNrp1 signaling in refinement, L2/3 PlexinD1 mutant neurons show abnormal heterotopic callosal projections to the contralateral striatum, possibly due to deregulated developmental refinement (Velona *et al.*, 2019).

In sum, we demonstrate that in the somatosensory cortex, Nrp1 regulates the developmental postnatal growth and refinement of L2/3 callosal axons. In this manner, the Nrp1 gradient determines balanced homotopic and heterotopic interhemispheric connectivity between primary and secondary somatosensory circuits.

## Methods

### Animals

Wild-type (WT) C57BL6JRccHsd (Envigo Laboratories, formerly Harlan. Indianapolis, the U.S.) mice were used in all experiments. The morning of the day of the appearance of a vaginal plug was defined as embryonic day 0.5 (E 0.5). Animals were housed and maintained following the guidelines from the European Union Council Directive (86/609/European Economic Community). All procedures for handling and sacrificing complied with all relevant ethical regulations for animal testing and research. All experiments were performed under the European Commission guidelines (2010/63/EU) and were approved by the CSIC and the Community of Madrid Ethics Committees on Animal Experimentation in compliance with national and European legislation (PROEX 124-17; 123-17).

### *In utero* electroporation and plasmids

Plasmids used were pCAG-GFP (Addgene, plasmid #11150), pCAG-Nrp1 (gift from Prof. Mu-ming Poo), and shNrp1 in pLKO.1 vector (hairpin sequence: CCTGCTTTCTTCTCTTGGTTTC. #TRCN0000029859, Merck. Darmstadt. Germany). *In utero* electroporation was performed as previously described (Briz *et al.*, 2017). Briefly, a mixture of the specified plasmids at a concentration of 1 μg/ μl each (pCAG-GFP or pCAG-Nrp1) or 0.6 μg/ μl (pLKO.1-shNrp1) was injected into the embryo’s left lateral ventricle using pulled glass micropipette. Five voltage pulses (38 mv, 50ms) were applied using external paddles oriented to target the somatosensory cortex. After birth, P2 GFP^+^ pups were selected and allowed to develop normally until P14 and P28. After sectioning, analyses were performed only in animals in which the electroporated area included both S1 and S2.

### CTB injections for retrograde labeling

Retrograde labeling from the CC and the cortical plate were performed by injecting subunit B of cholera toxin (CTB) conjugated to Alexa Fluor 555 (#C-34776, ThermoFisher Scientific. Massachusetts, the U.S.). Injections were performed in the CC, close to the midline, as previously reported (De Leon Reyes *et al.*, 2019), or in the cortical plate; in both cases, in the contralateral non-electroporated hemisphere (right hemisphere). Stereotaxic coordinates, injection volumes, and procedures for different developmental stages for injections in CC were performed as previously described (De Leon Reyes *et al.*, 2019). For cortical plate injections at P30, stereotaxic coordinates (anteroposterior (AP), mediolateral (ML), and dorsoventral (DV) axes from Bregma) were adjusted using the atlas of Paxinos (Paxinos and Franklin, 2004) and used as follow: S1/S2 injections (−1.34 mm AP; +3.7 mm ML; −0.4 ~ −0.5 mm DV) and, S2 injections (−1.34 mm AP; +3,7 mm ML; −0.7 ~ −0.8 mm DV); injecting 100 nL of CTB solution at 4 nl s^−1^. Animals were anesthetized during the surgical procedure with isoflurane/oxygen and placed on a stereotaxic apparatus (Harvard Apparatus. Massachusetts, the U.S.). CTB particles (diluted at 0.5% in phosphate-buffered saline (PBS)) were injected with a Drummond Nanoject II Auto-Nanoliter Injector using 30 mm pulled glass micropipettes (3000205A and 3000203G/X. Drummond Scientific Co. Pennsylvania, the U.S.). Mice were intrapericardially perfused with formalin two days after the surgery and brains were extracted and fixed overnight in formalin at 4°C. After fixation, brains were cryoprotected with 30% sucrose (#S0389. Merck. Darmstadt. Germany) and frozen in Tissue-Tek® O.C.T.™ Compound (#4583, Sakura Tissue-Tek. Tokyo. Japan).

### Immunohistochemistry

50 μm free-floating brain cryosections were used for immunofluorescence. Rabbit polyclonal anti-GFP (#A11122, Thermo Fisher Scientific. Invitrogen. Massachusetts, the U.S.) was used as primary antibody and goat anti-rabbit-Alexa 488 (#A11034, Thermo Fisher Scientific. Life Technologies. Massachusetts, the U.S.) as the secondary antibody. Nuclei were stained with 4’,6-diamidino-2-phenylindole (DAPI) (#D9542, Merck. Darmstadt. Germany).

### Confocal imaging and quantification

Confocal microscopy was performed using a TCS-SP5 (Leica. Wetzlar. Germany) Laser Scanning System on Leica DMI8 microscopes. Up to 50 m optical z-sections were obtained by taking 3.5 m serial sections with LAS AF v1.8 software (Leica. Wetzlar. Germany). Tilescan mosaic images were reconstructed with Leica LAS AF software. All images were acquired using a 512 × 512 scan format with a 20x objective.

For the acquisition and quantifications of the fluorescence signal (Rodriguez-Tornos *et al.*, 2016; Briz *et al.*, 2017), detectors were set to ensure equivalent threshold and signal-to-noise ratios between all samples. The maximum threshold signal was set by ensuring that no pixels were saturated. The threshold for background noise was determined using regions outside of the electroporated area (Rodriguez-Tornos *et al.*, 2016; Briz *et al.*, 2017). This approach ensures linearity between samples. Quantification of innervation was performed in tilescan images of electroporated (ipsilateral) and non-electroporated (contralateral) hemispheres. Different areas were measured delimitating manually ROIs, adjusting the threshold above the noise (making a binary image), and measuring the integrated density (using Fiji-ImageJ (Schindelin *et al.*, 2012)). Measures of contralateral ROIs were normalized to ipsilateral ones to avoid any differences in electroporation efficiency. Contralateral normalizations, without considering ipsilateral signal, were calculated to confirm the results. To quantify CC fasciculation, a midline ROI was selected to measure the fluorescence profile throughout ten equal distance bins. The different profiles were plotted to identify changes in dorsoventral routes.

Quantification of CTB^+^ over GFP^+^ cells in the primary (S1) and secondary (S2) somatosensory areas was performed on single plane confocal images from z-stacks (De Leon Reyes *et al.*, 2019). The proportions of CTB^+^ cells were calculated among randomly selected GFP^+^ cells in the ipsilateral (electroporated) hemisphere. For quantification of GFP^−^ populations, the proportions of CTB^+^ cells were calculated over randomly selected DAPI^+^ cells, excluding GFP^+^ cells. Functional areas of the adult mouse brain were identified using the atlas of Paxinos (Paxinos and Franklin, 2004).

### Statistical analysis

Sample size was determined to be adequate based on the magnitude and consistency of measurable differences between groups. Each experimental condition was carried out with a minimum of three biological replicates, a minimum of two sections from each brain, and included a minimum total number of 300 counted cells. During experiments, investigators were not blinded to the electroporation condition of animals. Results are expressed as the mean ± standard error of the mean (SEM). Results were compared using two-way ANOVA and one-way ANOVA with posthoc comparison with Tukey and Bonferroni’s tests. Statistical tests were performed using Prism 8 software (GraphPad Software. California, the U.S.).

## Article and author information

Author details:

**F. Martín-Fernández**

National Center of Biotechnology. Consejo Superior de Investigaciones Científicas. CNB-CSIC. Spain.

Contribution: conceptualization, formal analysis, validation, investigation, visualization, methodology, writing.

**C. García-Briz**

National Center of Biotechnology. Consejo Superior de Investigaciones Científicas. CNB-CSIC. Spain. Present address: Ministry of Consumer Affairs. Spain Govern.

Contribution: conceptualization, investigation, methodology.

**M. Nieto**

National Center of Biotechnology. Consejo Superior de Investigaciones Científicas. CNB-CSIC. Spain.

Contribution: conceptualization, writing.

## Acknowledgments

We are grateful to R. Gutierrez, A. Morales, S. Gutiérrez-Erlandsson, and A. Oña for technical assistance. J. García-Marqués, L.A. Weiss, N. S. de León, I. Varela, E. Marcos, and L. Bragg for critical reading and advice. F. Martín-Fernández holds an FPU fellowship from the Spanish MEFP, FPU15/02111. C. García-Briz was supported by a fellowship from the Spanish MICINN, FPI-BES-2012-056011. This work was funded by grants from the Ministerio de Ciencia, Innovación y Universidades/Agencia Estatal de Investigación/Fondo Europeo de Desarrollo Regional, European Union (SAF2017-83117-R and RED2018-102553T).

## Competing Interests

The authors declare no competing interests.

---

**Figure 1 – figure supplement 1.**
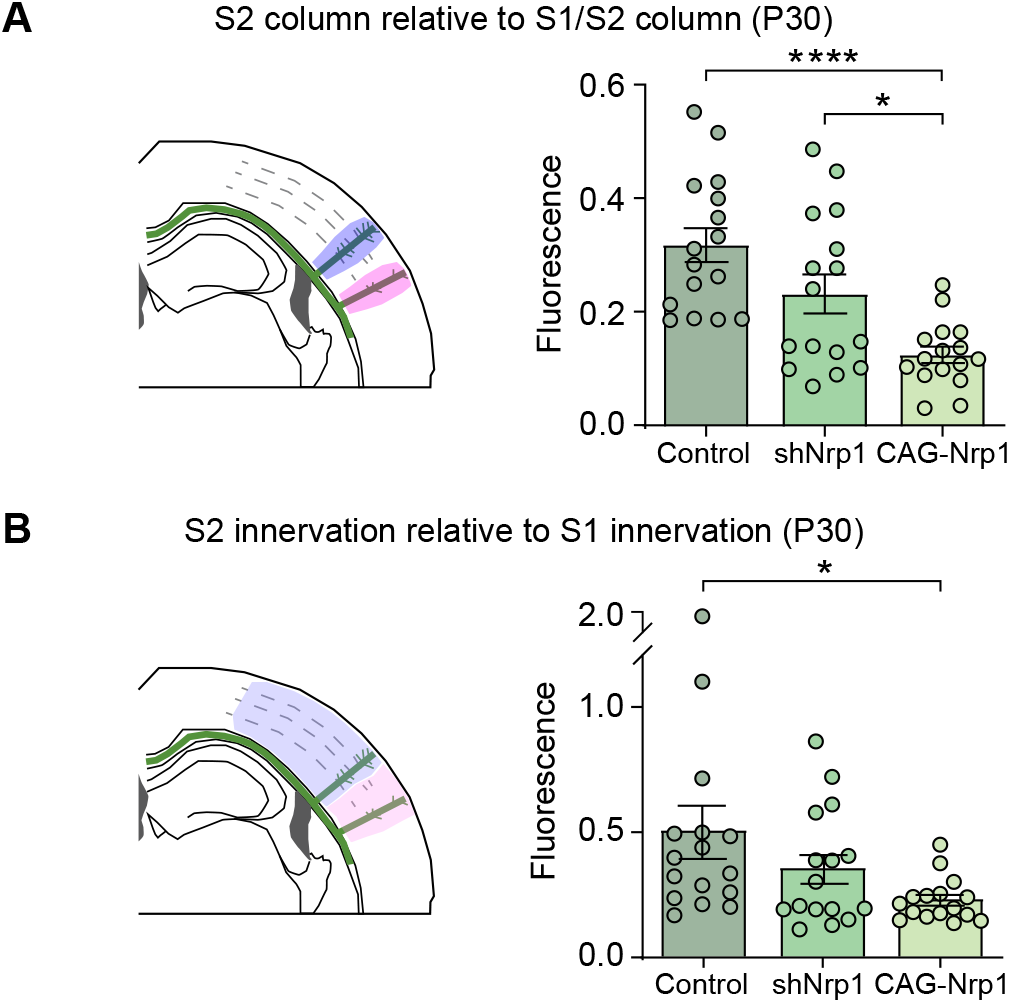
Analysis of contralateral innervation of SS cortex at P30 upon Nrp1 modifications. **A-B)** Quantification of axonal distribution in the contralateral hemisphere. The left panels depict schemes showing the selected ROIs in which GFP^+^ is quantified (shaded areas). Graphs show values of relative contralateral GFP innervation. Mean ± SEM (n = 8 brains, 2 sections per brain, in all conditions). A) S2 column relative to S1/S2 column (One-way ANOVA: *P*-value < 0.0001. Post-hoc with Tukey’s test: **** *p*-value _Control – CAG-Nrp1_ < 0.0001; * *p*-value _shNrp1 – CAG-Nrp1_ = 0.0231). B) S2 innervation relative to S1 innervation (One-way ANOVA: *P*-value = 0.0329 Posthoc with Tukey’s test: * *p*-value _Control – CAG-Nrp1_ = 0.0252).

**Figure 1 – figure supplement 2.**
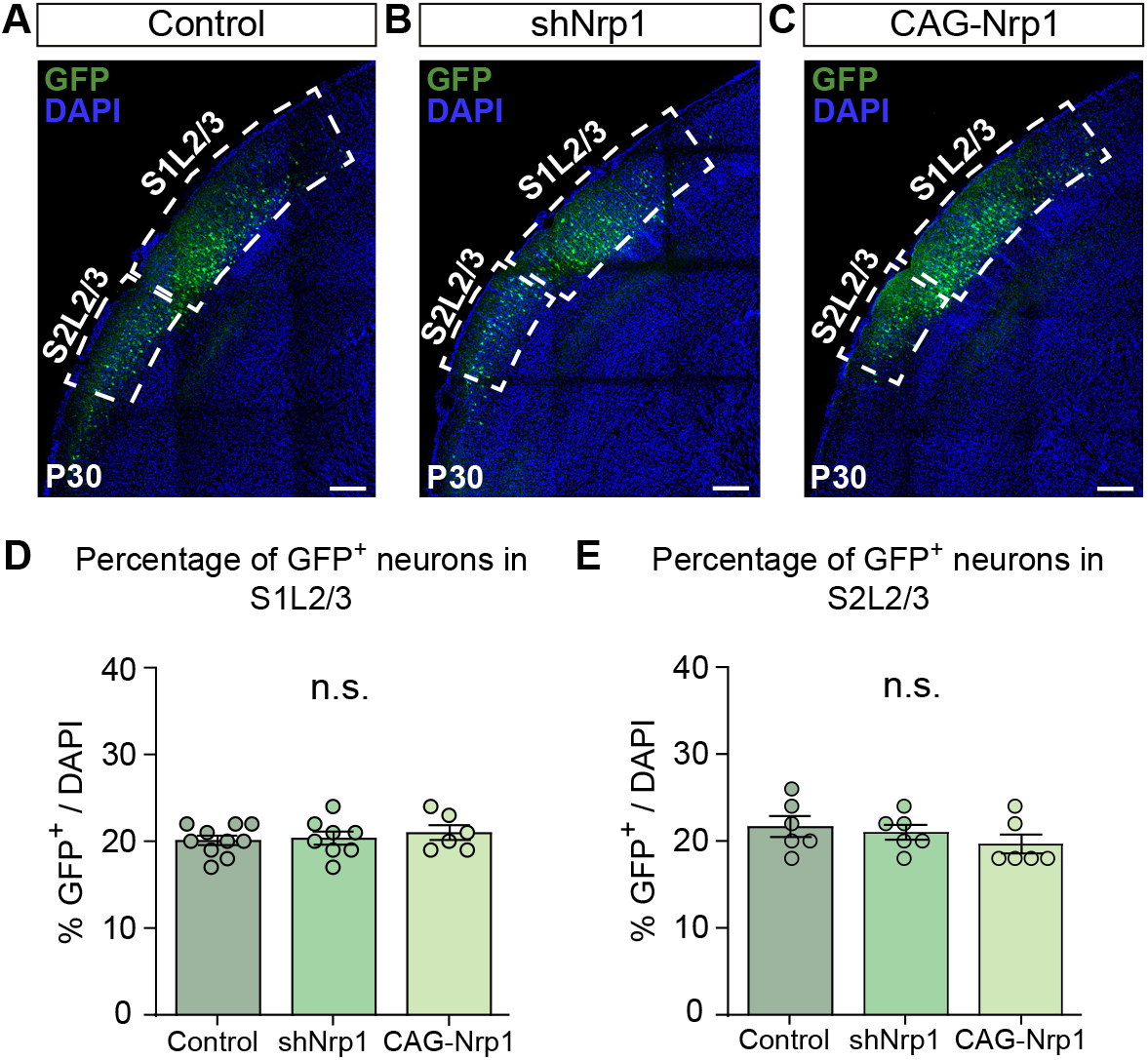
Analysis of proportions of GFP^+^ electroporated neurons in the ipsilateral side in somatosensory areas. **A-C)** Detail of the ipsilateral hemisphere of P30 electroporated brains from all conditions (Dashed boxes: S1L2/3 electroporated area and S2L2/3 electroporated area). Scale bar = 300 μm. **D-E)** Percentage of GFP^+^ neurons over cells (DAPI^+^) in S1L2/3 and S2L2/3. Mean ± SEM (n ≥ 3 brains, 2 sections per brain in all conditions). D) S1 area (One-way ANOVA: *P*-value = 0.6769 (n.s.)). E) S2 area (One-way ANOVA: *P*-value = 0.4172 (n.s.)).

**Figure 2 – figure supplement 1.**
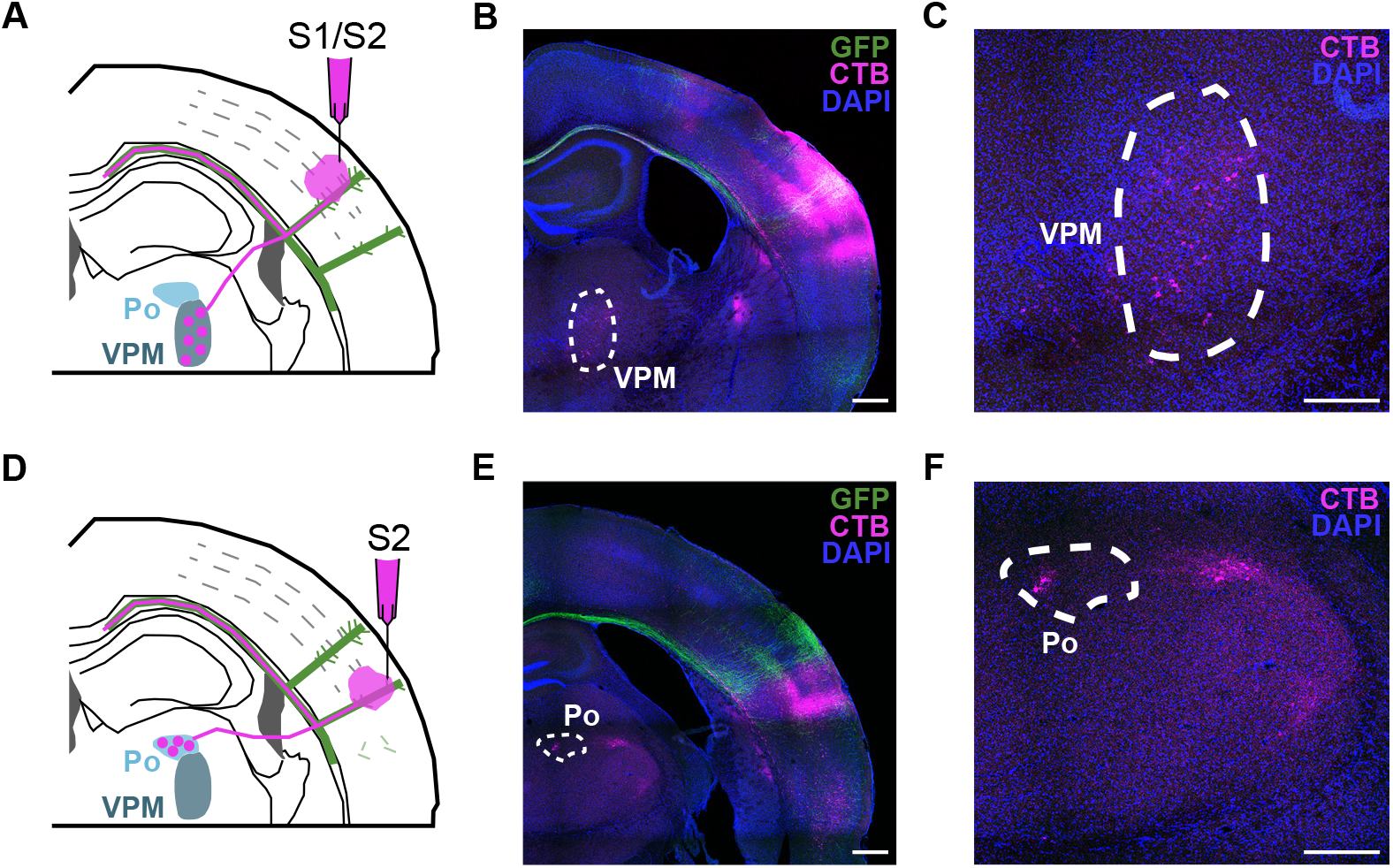
Retrospective control of correct columnar stereotaxic injections. **A)** Scheme of coronal section of P30 injected brain in S1/S2 column. Thalamus nucleus VPM (ventral posterio medial nuclei) extend thalamocortical axons to S1/S2 column. **B)** Merge image of coronal section of the contralateral injected side of S1/S2 injection. Dashed line marks VPM, where thalamic neurons are CTB^+^. Green = GFP, Magenta = CTB, Blue = DAPI. Scale bar = 500 μm. **C)** Detail of VPM nucleus. Magenta = CTB, Blue = DAPI. Scale bar = 200 μm. **D)** Scheme of coronal section of P30 injected brain in S2 column. Thalamus nucleus Po (posterior nucleus) extend thalamocortical axons to the S2 column. **E)** Merge image of coronal section of contralateral injected side of S2 injection. Dashed line marks Po, where thalamic neurons are CTB^+^. Green = GFP, Magenta = CTB, Blue = DAPI. Scale bar = 500 μm. **F)** Detail of Po nucleus. Magenta = CTB, Blue = DAPI. Scale bar = 200 μm.

**Figure 2 – figure supplement 2.**
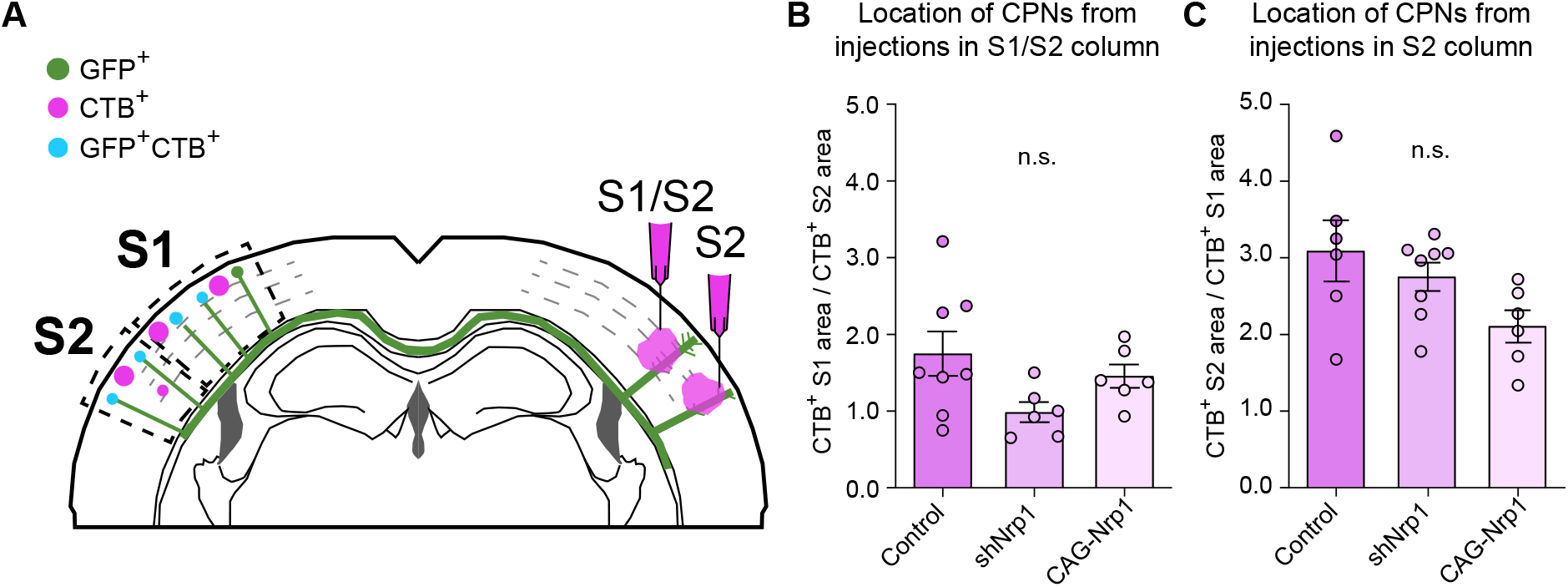
Analysis of location of CPNs neurons at P30 in somatosensory cortex. **A)** Scheme of coronal section of electroporated and injected brain at P30. The injections were made in S1/S2 column in one group and in S2 column in the other group. CTB^+^ and GFP^+^CTB^+^ populations were quantified individually between the S1 and S2 area and, in the two types of injections. **B-C)** Location of CTB^+^ CPNs from injections in S1/S2 column and S2 column. For S1/S2 injections, ratio was calculated between S1CTB^+^ and S2CTB^+^ neurons. For S2 injections, ratio was calculated between S2CTB^+^ and S1CTB^+^ neurons. Mean ± SEM (n ≥ 3 brains, 2 sections per brain in all conditions. B) S1/S2 injections (One-way ANOVA: *P*-value = 0.0840 (n.s.)). C) S2 injections (One-way ANOVA: *P*-value = 0.0669 (n.s.)).

**Figure 3 – figure supplement 1.**
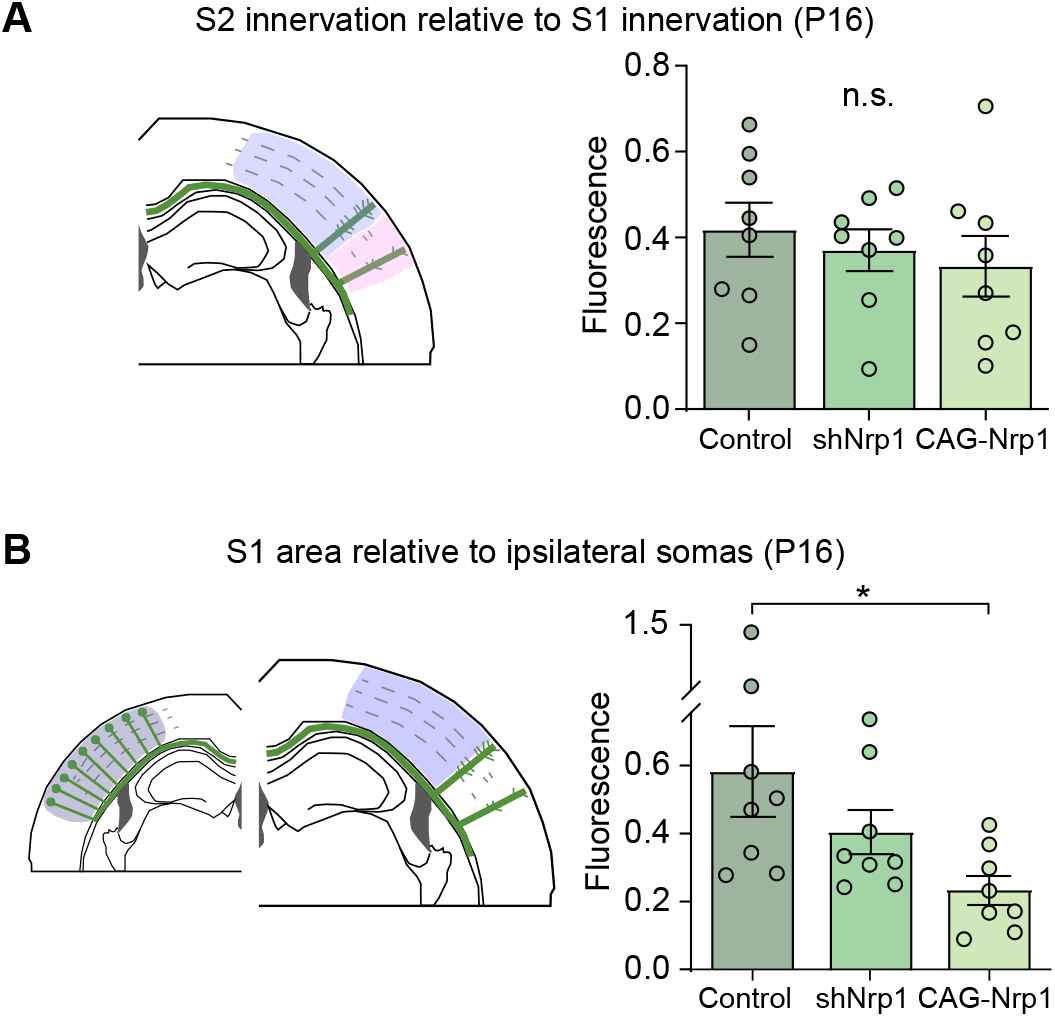
Analysis of contralateral innervation of SS cortex at P16 upon Nrp1 modifications. **A-B)** Quantification of contralateral GFP^+^ axons. Mean ± SEM (n ≥ 4 brains, 2 sections per brain in all conditions). A) S2 innervation relative to S1 innervation (One-way ANOVA: *P*-value = 0.6260 (n.s.)). B) S1 area (One-way ANOVA: *P*-value = 0.0385. Posthoc with Tukey’s test: * *p*-value _Control – CAG-Nrp1_ = 0.0300).

**Figure 5 – figure supplement 1.**
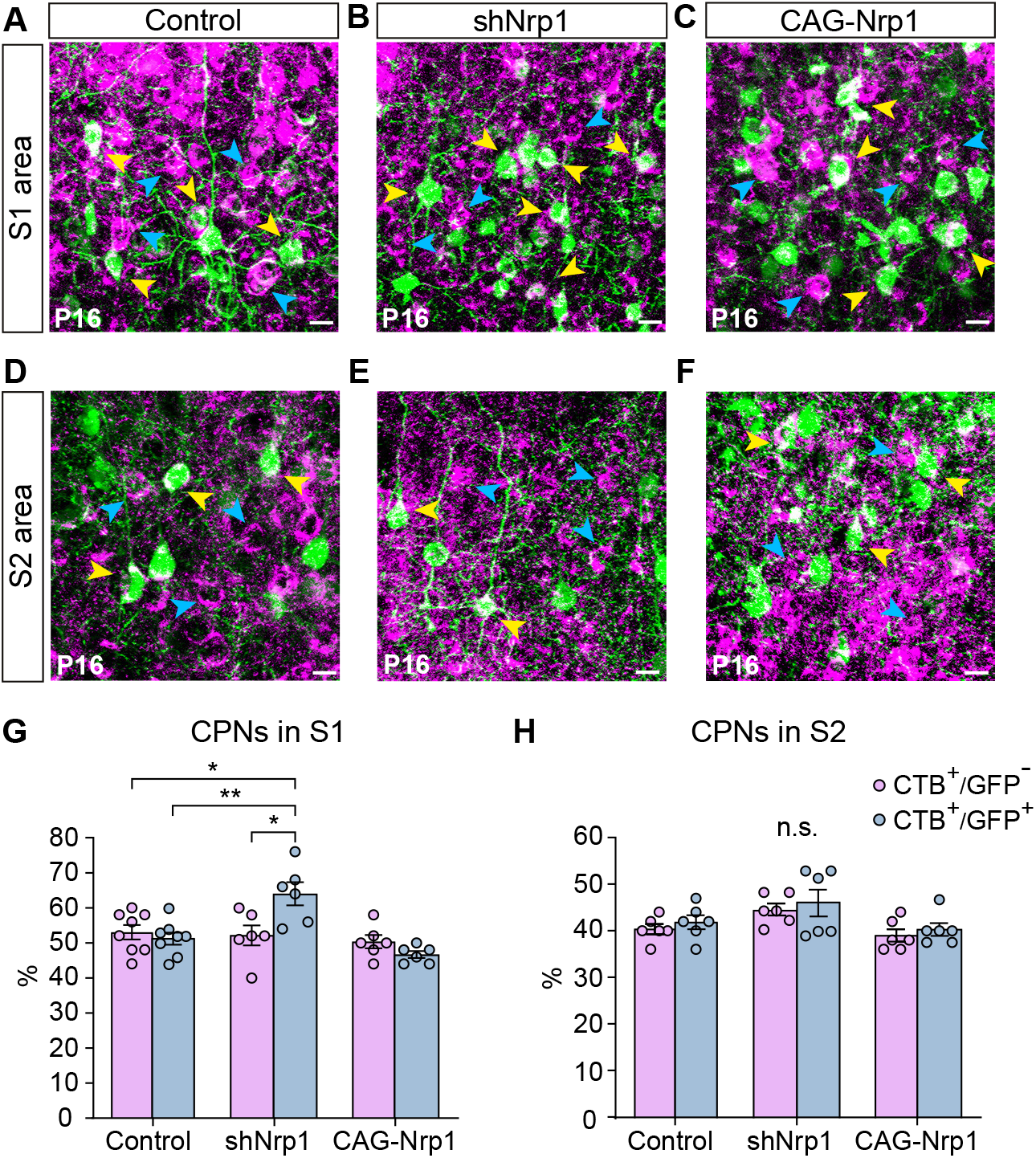
CPNs proportions of non-electroporated and electroporated neurons at P16. **A-F)** Images of L2/3 populations (CTB^+^, blue arrowheads; GFP^+^CTB^+^, yellow arrowheads). Scale bar = 10 μm. A-C) S1L2/3 neurons. D-F) S2L2/3 neurons. **G-H)** Proportion of CTB^+^/GFP^−^ neurons, and GFP^+^CTB^+^/GFP^+^ in S1 area and S2 area. Mean ± SEM (n ≥ 3 brains, 2 sections per brain in all conditions). G) S1 CPNs (Two-way ANOVA: *P*-value _Experimental condition_ = 0.001; *P*-value _Population_ = 0.242. Posthoc with Tukey’s test: * *p*-value _shNrp1 GFP- – shNrp1 GFP+_ = 0.0127; ** *p*-value _Control GFP+ – shNrp1 GFP+_ = 0.0033; * *p*-value _Control GFP- - shNrp1 GFP+_ = 0.0135). H) S2 CPNs (Two-way ANOVA: *P*-value _Experimental condition_ = 0.072 (n.s.); *P*-value _Population_ = 0.2842).

**Figure 5 – figure supplement 2.**
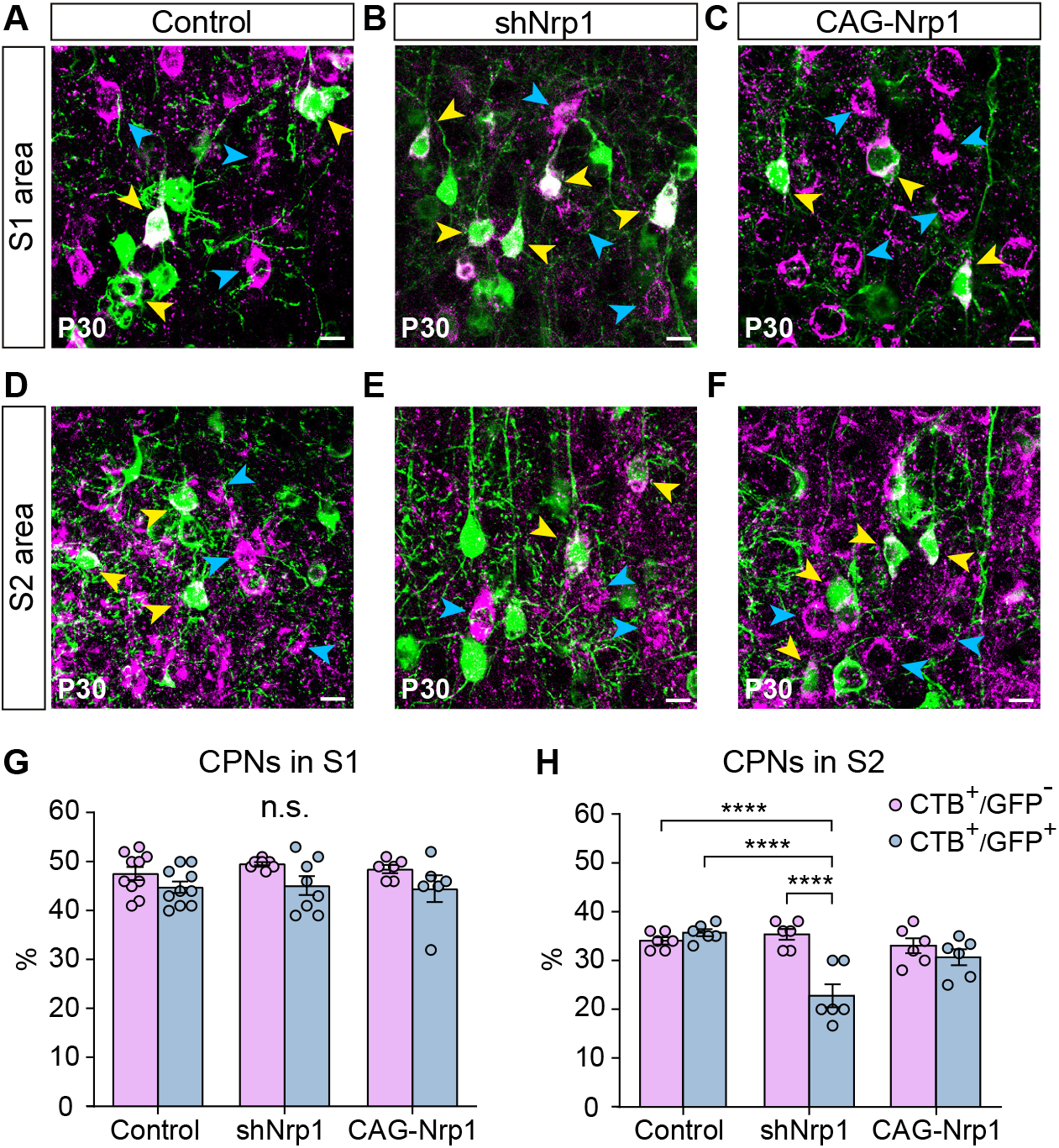
CPNs proportions of non-electroporated and electroporated neurons at P30. **A-F)** Images of L2/3 populations (CTB^+^, blue arrowheads; GFP^+^CTB^+^, yellow arrowheads). Scale bar = 10 μm. A-C) S1L2/3 neurons. D-F) S2L2/3 neurons. **G-H)** Proportion of CTB^+^/GFP^−^ neurons, and GFP^+^CT-B^+^/GFP^+^ in S1 area and S2 area. Mean ± SEM (n ≥ 3 brains, 2 sections per brain in all conditions). G) S1 CPNs (Two-way ANOVA: *P*-value _Experimental condition_ = 0.6998 (n.s.); *P*-value _Population_ = 0.0041). H) S2 CPNs (Two-way ANOVA: *P*-value _Experimental condition_ = 0.0018; *P*-value _Population_ = 0.0008. Posthoc with Tukey’s test: **** *p*-value _shNrp1 CTB+GFP- – shNrp1 CTB+GFP+_ < 0.0001; **** *p*-value _Control CTB+GFP+ – shNrp1 CTB+GFP+_ < 0.0001; **** *p*-value Control _CTB+GFP- – shNrp1 CTB+GFP+_ < 0.0001).

## Notes

### Competing Interest Statement

The authors have declared no competing interest.

